# Hierarchical Bayesian Models of Reinforcement Learning: Introduction and comparison to alternative methods

**DOI:** 10.1101/2020.10.19.345512

**Authors:** Camilla van Geen, Raphael T. Gerraty

## Abstract

Reinforcement learning models have been used extensively to capture learning and decision-making processes in humans and other organisms. One essential goal of these computational models is the generalization to new sets of observations. Extracting parameters that can reliably predict out-of-sample data can be difficult, however. The use of prior distributions to regularize parameter estimates has been shown to help remedy this issue. While previous research has suggested that empirical priors estimated from a separate dataset improve predictive accuracy, this paper outlines an alternate method for the derivation of empirical priors: hierarchical Bayesian modeling. We provide a detailed introduction to this method, and show that using hierarchical models to simultaneously extract and impose empirical priors leads to better out-of-sample prediction while being more data efficient.

## Introduction

First-hand experience and decades of behavioral studies teach us that reward is one of the most powerful drivers of learning, and that behavior is often modified to maximize its acquisition (Gormezano & Moore, 1966; Rescorla & Wagner, 1972; Schultz et al., 1997; Skinner, 1935; Sutton et al., 1991; Thorndike, 1898; Yerkes & Morgulis, 1909). Although our daily lives are ripe with examples of this phenomenon, the question of how to accurately quantify reward’s influence on our actions remains an active area of research. A quantitative theory of reward learning is crucial; it provides a necessary bridge towards understanding the internal computations underlying behavior.

One popular framework for modeling learning and decision-making processes is reinforcement learning (RL) (Daw & Doya, 2006; Niv, 2009; Sutton & Barto, 1990, 1998). A pervasive issue with reinforcement learning models, however, is how to handle variability between people (Ballard & McClure, 2019; Gershman, 2016; Katahira, 2016; Shahar et al., 2019). In this paper, we provide an introduction to one emerging set of strategies for handling this variability: hierarchical Bayesian models. Our goals are to 1) give a detailed description of hierarchical models and their application in the context of reinforcement learning and 2) compare these models to other commonly used approaches. We show that on a simple but widely used reinforcement learning task, hierarchical Bayesian models provide the best predictive accuracy compared to other methods, including a recently suggested technique that relies on the collection of a separate dataset to enable robust inference (Gershman, 2016).

Given the many different uses of the term reinforcement learning, we must first specify the scope of this paper. In its broadest sense, RL refers to a set of problems and solutions centered on learning from interacting with an environment without explicit supervision. In neuroscience and psychology, RL is usually used to refer to a subset of RL algorithms that are used to model neural and behavioral data. In this review, we will narrow the scope even further, to those RL models that describe how humans (and other organisms) learn behavioral policies from rewards (Daw & Doya, 2006; Niv, 2009; Sutton & Barto, 1990, 1998). In their most basic formulations, these RL models estimate the probability of a specific action in a given context based on two determining factors: that action’s value in that context (learned through its reward history), and how influential this value will be in determining choice. When fitting these models to data from humans or other organisms, both of these factors can vary from one individual to the next. To manage this variability, standard RL models fit two individual-specific parameters: learning rate and inverse temperature. The learning rate (*α*) determines the extent to which prediction error will play a role in updating an action’s value. This prediction error is quantified as the difference between the expected value of an action and the actual outcome on a given trial. Higher values of *α* imply greater sensitivity to the most recent choice outcome, while lower *α*s are indicative of more gradual value updating. Once value is computed, people can also differ in how much influence it exerts on their behavior, or how exploratory they are in their choices. This tendency is governed by an inverse temperature parameter (*β*) whose magnitude determines the impact of value on choice. We will provide more precise mathematical formulations of RL models below, but to begin with, it is worth considering the overarching goal of fitting them – or of fitting any statistical model – in the first place.

In using RL models to summarize or compress observed data, we seek to extract meaningful structure from these observations. To this end, we want our models to generalize beyond the specific data we use to fit them. With this goal in mind, a common way of evaluating models is out-of-sample prediction (Akaike, 1998; Arlot & Celisse, 2010; Vehtari et al., 2017). In RL models, for instance, we might ask how well our parameters for any specific subject predict future data from that individual, or how well our group-level averages predict data from new subjects (Daw, 2011).

Extracting parameter values that can reliably predict out-of-sample data is not a trivial task, however. When fitting computational models, we must strike a balance between describing the real patterns in the data without mistakenly inferring patterns from noise. In the context of RL models, different combinations of parameter values can sometimes fit a subject’s data similarly well. If this is the case, differences between these plausible sets of parameters will in part be due to noise, and are unlikely to replicate out-of-sample. Thus, without further constraints, these RL models are likely to lead to poor out-of-sample predictive accuracy, and the parameter estimates derived from them are likely to be overconfident.

One way to counter the overconfidence that results from selecting just one set of estimates has to do with the distinction between modes (or maxima) and expectations over full distributions. While maximum likelihood or maximum a posteriori methods use the parameter values with the best fit (based on the mode of the likelihood or posterior distribution), a more fully Bayesian approach is to approximate the posterior distribution of all possible values, and take the expectation of any functions of the parameters over the full distribution. For example, we might be interested in the correlation between parameters fit to multiple subjects and some other measure. If we are working in a Bayesian setting, we might compute the expectation of this correlation over the posterior, or even generate a posterior distribution over correlation values. In addition, we are often interested in the marginal posterior distribution of one or two of the model’s parameters, which is obtained by integrating the joint posterior distribution over all other parameters. For example, we might be interested in a single individual’s learning rate, which would have a different posterior distribution for different inverse temperature values. Rather than selecting a point estimate based on the most likely value, the marginal posterior of learning rate would be obtained by integrating the posterior distribution over inverse temperatures (put differently, it is the expectation of p(*α*|*β*, Y) taken over p(*β*|Y), with Y corresponding to the choice data).

In some simple cases, the Bayesian and the maximum likelihood approach will lead to identical results. However, when fitting models with hierarchical structure, many dimensions, or parameters whose more likely values may be close to boundaries (such as variance, which cannot be less than zero), these two summary statistics can differ substantially. In this paper, we will not directly compare modal vs. Bayesian methods, but for more direct demonstrations of the benefits of the Bayesian approach, see Asparouhov & Muthén, 2020; Browne & Draper, 2006; Gelman et al., 2013.

Within the Bayesian framework, there are still many ways to approximate posterior distributions given some data and a model. One popular choice is Markov Chain Monte Carlo (MCMC), which has the benefit of not requiring assumptions about the shape of the posterior distribution, and is the method we will use in the subsequent sections. In brief, MCMC is a method that approximates the posterior distribution by iteratively taking samples from it. For a sufficiently high sample size, MCMC approximations will converge to the true posterior distribution, though convergence checks are necessary to verify that this is the case. Additional resources concerning convergence checks for MCMC sampling can be found in the “bayesplot” package in Stan, but are beyond the scope of this tutorial (https://mc-stan.org/bayesplot/articles/visual-mcmc-diagnostics.html).

Another commonly used strategy to improve predictive accuracy is regularization: using a bias term to penalize overfitting. While regularization is not inherently Bayesian, regularization penalties can often be expressed as prior distributions (Cao & Ray, 2012; Efron, 1996). In general, these prior distributions constrain the values of model parameters by biasing estimates towards regions of parameter space considered more plausible a priori. By setting priors on parameter values, we are inserting inductive bias into the model in the hopes of minimizing the variance of our estimates over repeated samples, which can translate to greater predictive accuracy (Briscoe & Feldman, 2006). Given the tradeoff between fit to a specific dataset and generalizability, the inclusion of priors is one way of balancing the two extremes (Gershman, 2016). However, picking too strong a prior may mean that we overly bias estimates and miss patterns in the data, whereas a weak prior might lead to overfitting. Thus, in addition to whether or not to include priors in RL models, the best practice for doing so remains an open question.

For data with the hierarchical or repeated-measures structure often found in psychology and neuroscience experiments, one way to generate priors for lower-level observations is to extract them empirically from group-level data. The parameter estimates for individual subjects in an experiment, for example, would be biased towards a group mean based on a population distribution of subject-level parameters. This principle of utilizing group-level data to inform individual-level estimates underlies mixed-effects modeling (Baayen et al., 2008; Barr, 2013; Bates, 2005), which can use the maximum likelihood approach described above, as well as the fully Bayesian hierarchical models we will describe here.

In the case of RL models, it has recently been suggested that empirical priors can be estimated from the group distributions derived from a separate dataset (Gershman, 2016). Doing so constrains individual variability in parameter estimates based on the behavior of a separate group of participants on the same task. Although this method has been shown to improve predictive performance, it requires a large dataset to draw from: a substantial subset of subjects is used to generate group-level priors and then discarded from the final model. In this paper, we will demonstrate that in a simple choice task, a hierarchical Bayesian approach to fitting reinforcement learning models actually predicts new data points better, while being more data efficient. In the next section, we will provide a detailed overview of the hierarchical model’s implementation. We will then use this approach to compare different reinforcement learning models and finally compare the hierarchical Bayesian approach to other ways of modeling the data, including the two-dataset approach described above.

Several neuroscience and psychology papers have already implemented the hierarchical Bayesian approach to fitting reinforcement learning models (Fontanesi et al., 2019; Guitart-Masip et al., 2012; Q. Huys et al., 2011; Schaaf et al., 2019; A. Wilson et al., 2007) and others have described its benefits explicitly (Katahira, 2016). Our goal here is to provide an introductory overview of these methods, as well as a quantitative evaluation of their generalization performance compared to other commonly used techniques.

## Model (M1)

To illustrate the hierarchical Bayesian approach, we first fit a standard computational model for so-called ‘bandit’ problems, where an individual makes repeated choices in the same environmental context or *state* (Sutton & Barto, 1998; Daw, 2011). We used the data from Gershman (2015) – available at https://github.com/sjgershm/RL-models – to fit this model. The dataset consists of choice behavior from 205 participants pooled across five different studies, each consisting of four 25-trial blocks. In all five studies, participants were instructed to choose one of two colored buttons, based on which one they believed had a higher expected reward. A common RL approach to this type of problem is to fit a simplified version of a Q-learning model to the data. In this model, the value of each of the two options was initialized at 0.5 and updated after each trial as the sum of the predicted value of the chosen option 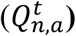 and a reward prediction error 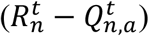, weighted by *α*_*n*_. More formally, this update corresponds to:

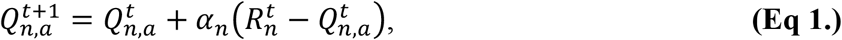

where *n* refers to subject, *t* to trial, and *a* to left or right choice. The likelihood of choosing left or right is then modeled as a Bernoulli distribution governed by parameter 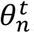, where 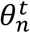 is a softmax transformation of the action values, weighted by a subject-specific *β*_*n*,1_. Because there are only two choice options, the softmax simplifies to a logistic transform. Higher values of *β*_*n*,1_ correspond to a greater bias towards the option that has a higher estimated value. Because participants used both hands to make button presses and given the prevalence of right-handedness in the population, we also included an intercept term (*β*_*n*,0_) to account for bias towards pressing one side more than the other. Characterizing choice 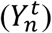 as either 0 (choose right) or 1 (choose left), we can model the choice probability as:

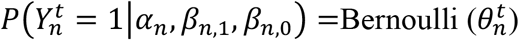

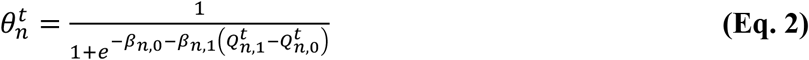

This represents the likelihood of an observed choice according to our model, given a set of parameter values.

However, we have yet to address how to most reliably extract these parameters from the data. As we mentioned above, fitting unbiased estimates maximizes the likelihood of each participants’ data, but might lead to lower predictive accuracy. In order to address this challenge, we include empirical priors on *α*_*n*_, *β*_*n*,1_ and *β*_*n*,0_ according to the distributions listed below, which are models of how the parameters are distributed in the population we have sampled from. For computational efficiency, *β*_*n*_ is a vector of *β*_*n*,1_ and *β*_*n*0_ parameters for a single participant.

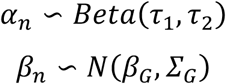

Here, *τ*_1_ and *τ*_2_ are shape parameters that describe the distribution of *α* across subjects, and thus constrain the subject-level estimates, which are assumed to follow a Beta distribution. Similarly, the subject-level *β*_*n*_ vectors are assumed to follow a multivariate normal distribution, where *β*_*G*_ is a vector of the population means for the slope and intercept. *Σ*_*G*_ is a matrix that includes the variance around the group-level distribution of the slope 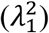, the variance around the group-level distribution of the intercept 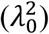, and the covariance between the two parameters across subjects. The magnitude of 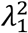 and 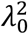 tell us the extent to which individuals differ in their intercept (right bias) and slope (inverse temperature), respectively. The larger the variances, the more we allow individual parameter values to stray from the group means. Similarly, the inter-individual variance for learning rate can be computed as the variance of the Beta distribution specified by *τ*_1_ and *τ*_2_ as follows:

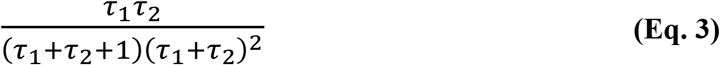

In addition to regularizing the subject-level parameters based on group-level data, we will also set weakly informative hyperpriors on the parameters of the group distributions themselves. These hyperprior distributions are chosen for computational convenience and to have little impact on inference. The heavy-tailed nature of the Cauchy distribution helps ensure that the hyperpriors will be only weakly informative, and we use a positive half-Cauchy to guarantee that the parameter values will be positive (as is necessary to specify a Beta distribution). In other settings (strong prior knowledge about the group distributions or different estimation methods), other hyperpriors may make more sense. Here, they are defined according to the following distributions:

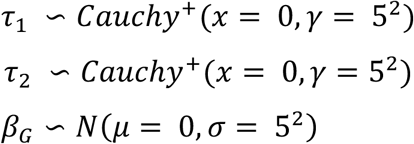

We will also set a hyperprior on *Σ*_*G*_. It can be more convenient to specify prior distributions on covariance matrices in terms of correlation matrices and standard deviations. This is the case because unlike variances and covariances, standard deviations and correlations are, respectively, on the same scale as the parameters and scale-free. This makes it easier for us to have an intuitive sense of how they are operating. Furthermore, we might have more prior information about the standard deviation/variance than we do about the covariance. In this case, it is useful to be able to state separate, more informative priors for the standard deviations and less informative priors on the correlations. Though it is beyond the scope of this paper, Barnard, McCulloch, et al., 2000 provide a detailed explanation of these and other issues relating to modeling covariance matrices.

In order to parameterize covariance in terms of correlations and standard deviations, we first construct two matrices with standard deviations *λ*_0_ and *λ*_1_ along their diagonals. We will then multiply these by a correlation matrix (*Ω*) as follows (Barnard et al., 2000; Gelman & Hill, 2007):

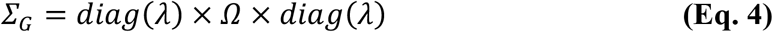

Thus, the between-subject covariance for *β* parameters is parameterized through the standard deviations of and correlations between those parameters. Hyperpriors can then be set on the components of *Σ*_*G*_ such that:

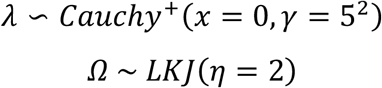

Here, we use the LKJ distribution, following Lewandowski et al., 2009. It is a distribution for sampling random correlation matrices, with a single parameter governing the distribution of correlation values. As this parameter gets larger, the samples become increasingly close to identity matrices, meaning zero correlation.

Overall, the parameters of the hierarchical model relate to one another as illustrated in the graphical depiction in Figure 1. Hyperpriors constrain group-level estimates, which constrain subject-level parameters that describe an individual’s choices.

**Figure 1:**
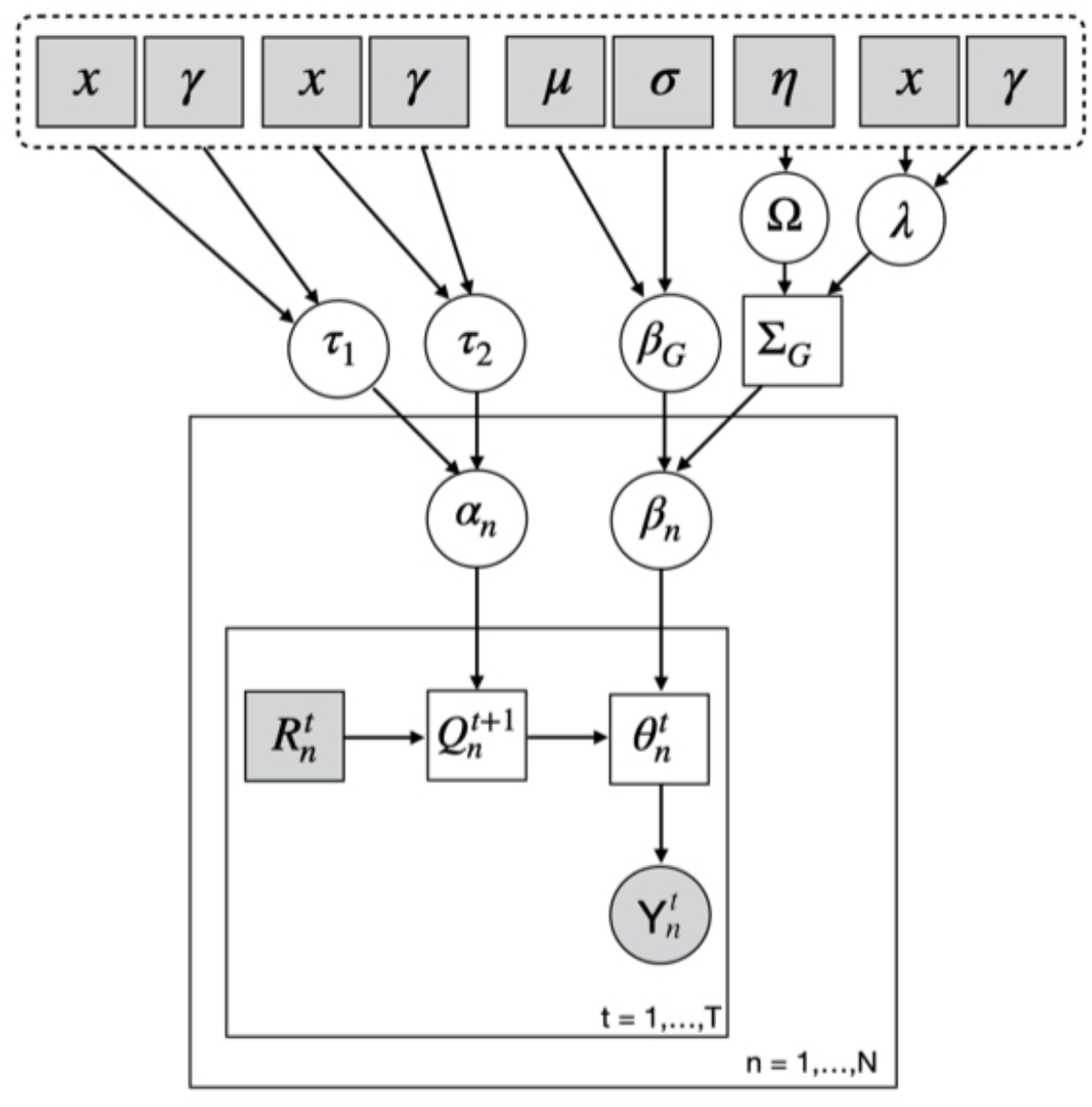
Graphical depiction of a hierarchical Bayesian model of standard Q-learning. Dashed line delineates the hyperpriors, which are set according to the following specifications in the text: *x* = 0, *γ* = 5; *μ* = 0, *σ* = 5; *η* = 2. Arrows are used to indicate interdependencies between variables. Squares denote deterministic variables, while random variables are illustrated by circular nodes. Observed variables are shaded in grey, while unobserved variables are white. *θ* corresponds to the likelihood of an observed choice, according to the model.

Finally, we fit the reinforcement learning model using Hamiltonian Markov Chain Monte Carlo in Stan (Carpenter et al., 2017) with four chains of two thousand iterations (including warmup). MCMC methods allow us to sample arbitrary posterior distributions by iteratively deciding whether the current sample is better or worse than the previous sample. As a result of this process, we will ultimately end up having approximated the true posterior distribution with our samples (Betancourt & Girolami, 2013). In the hierarchical setting, MCMC sampling allows us to estimate the posterior distribution over all subject- and group-level parameters simultaneously.

The marginal posterior distributions for the parameters of the group-level variables in the model described above can be found in Figure 2A. The empirical prior distributions that result from the expectations of these parameters are illustrated in Figure 2B. Both the group-level posteriors and the resulting priors are in line with previously reported results (Davidow et al., 2016; Daw et al., 2011; Eckstein et al., 2020; O’Doherty et al., 2007). Empirical priors have the effect of moving uncertain outlier estimates closer to the group prior, a phenomenon referred to as shrinkage. An illustration of the parameter shrinkage that results from its use of empirical priors for regularization can be found in Supplemental Figure 1.

**Figure 2:**
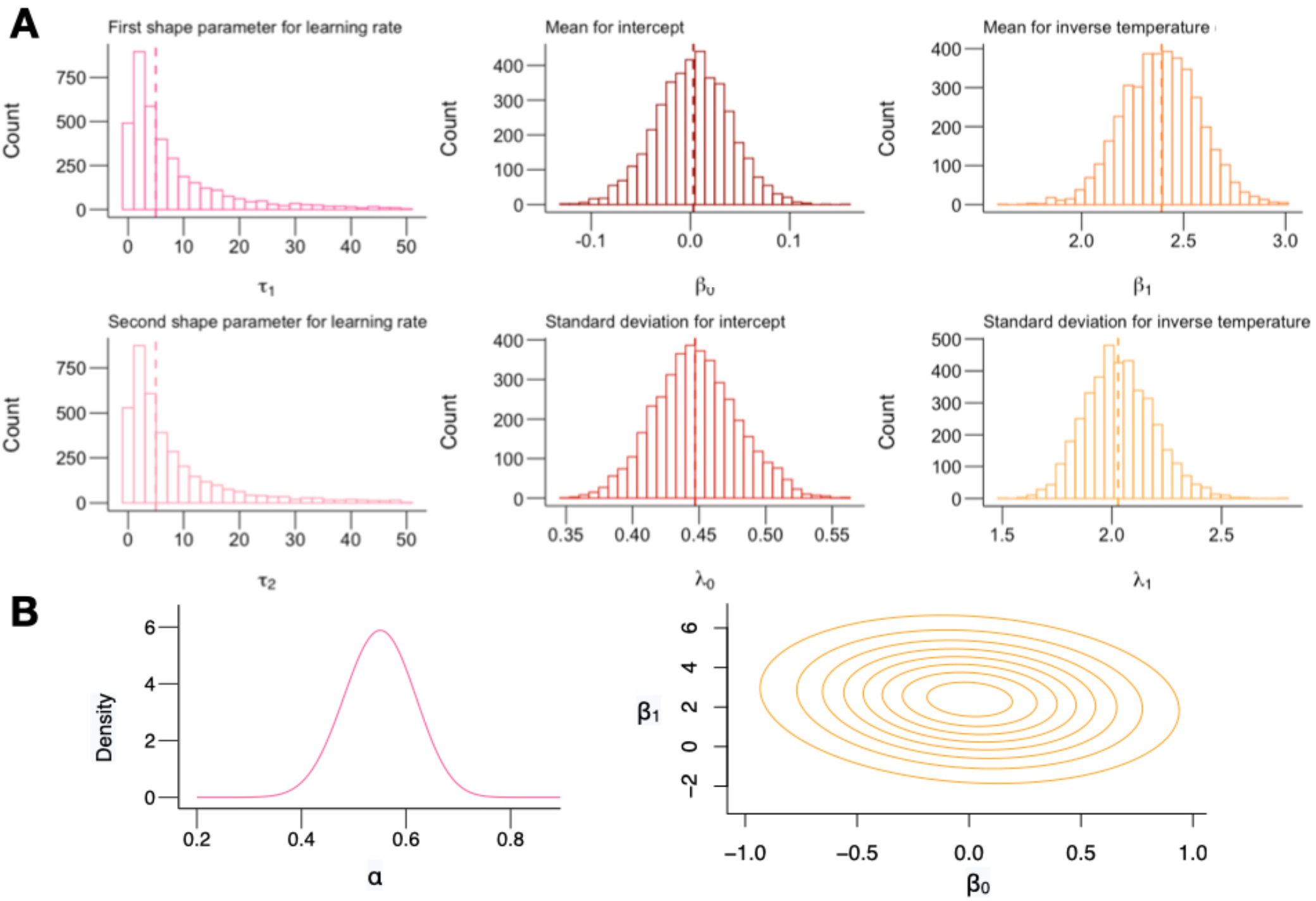
Posterior histograms for group-level parameters and expected empirical prior distributions in model M1. **(A)** Posterior histograms for group-level parameter estimates fit hierarchically on model M1. Dashed line corresponds to the median of the each distribution. The distributions for *τ*_1_and *τ*_2_ have been truncated at 50 for legibility. **(B)** Empirical prior distributions for α, β_0_ and β_1_. The density functions are parametrized using the mean of each parameter’s posterior distribution (from A). As specified by the model, the empirical prior on α follows a Beta distribution, while the empirical prior on β_0_. and β_1_ is multivariate Normal (the posterior distribution over the covariance is omitted from A).

In the next section, we will evaluate the simple Q-learning model’s performance in comparison to two more complex models. We will then use the best fitting reinforcement learning model to compare the predictive accuracy of the hierarchical Bayesian approach to that of three commonly used alternatives: one that allows for no subject-level variability and only fits the model at the level of the group, one that allows for infinite subject-level variability, which is equivalent to fitting a separate model for every subject, and another in which group-level empirical priors are extracted from a subset of the data and then fit on the remaining group (Gershman, 2016). We show that the hierarchical approach outperforms all of these alternatives in terms of accurately predicting new data.

### Model Comparison

The model we have described so far (M1) is a standard Q-learning model with one learning rate. Following the method used in Gershman 2016, we compared its performance to two alternate models with additional parameters:

‐ **M2**: Dual learning rates. This model uses the same choice function and the same set of priors as M1, but allows for two different learning rates depending on whether the prediction error was positive or negative. In other words, if 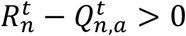, the Q-value is updated such that:

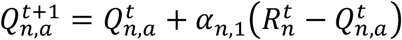

If 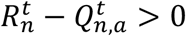, however, then the Q-value is updated such that:

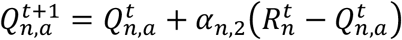

This two-learning-rate model is commonly used in the literature, and allows for differential value-updating mechanisms for outcomes that are better or worse than expected (Daw et al., 2002; Frank et al., 2009; Gershman, 2015; Niv et al., 2012).
‐ **M3**: Dual learning rates + stickiness. This model is identical to M2 but includes an additional stickiness variable *ω* that captures participants’ tendency to repeat the same choice as on the previous trial, regardless of whether it was rewarded. In practice, adding this parameter consists of updating the choice function such that:

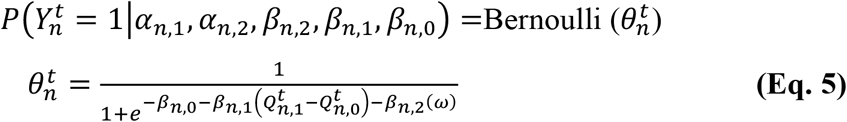

where *ω* takes on a value of 0.5 when the previous choice was left, and a value of -0.5 when the previous choice was right. With the added stickiness parameter, the model allows for 3 group-level and subject-specific *β* parameters, which are constrained according to the same priors as denoted above. The only change is that since there are now 3 group-level *β* parameters and the covariance matrix for our group-level *β* distribution is 3×3.

To compare the three models of interest – single learning rate (M1), dual learning rate (M2), and dual learning rate plus stickiness (M3) – we fit each reinforcement learning model using the MCMC method described above. To evaluate the fit of each model, we fit all of them on 3 out of 4 experimental blocks for each participant and computed the average log-likelihood of observations in the held-out block. The results are reported as the average log-likelihood for an observation for each participant, averaged across trials and posterior samples. As shown in Figure 3A and C, the model with dual learning rates and stickiness (M3) results in higher expected log likelihood for the held-out data (t(204) = 2.76; mean = 0.063 [0.018; 0.11]; p = 0.0063).

**Figure 3:**
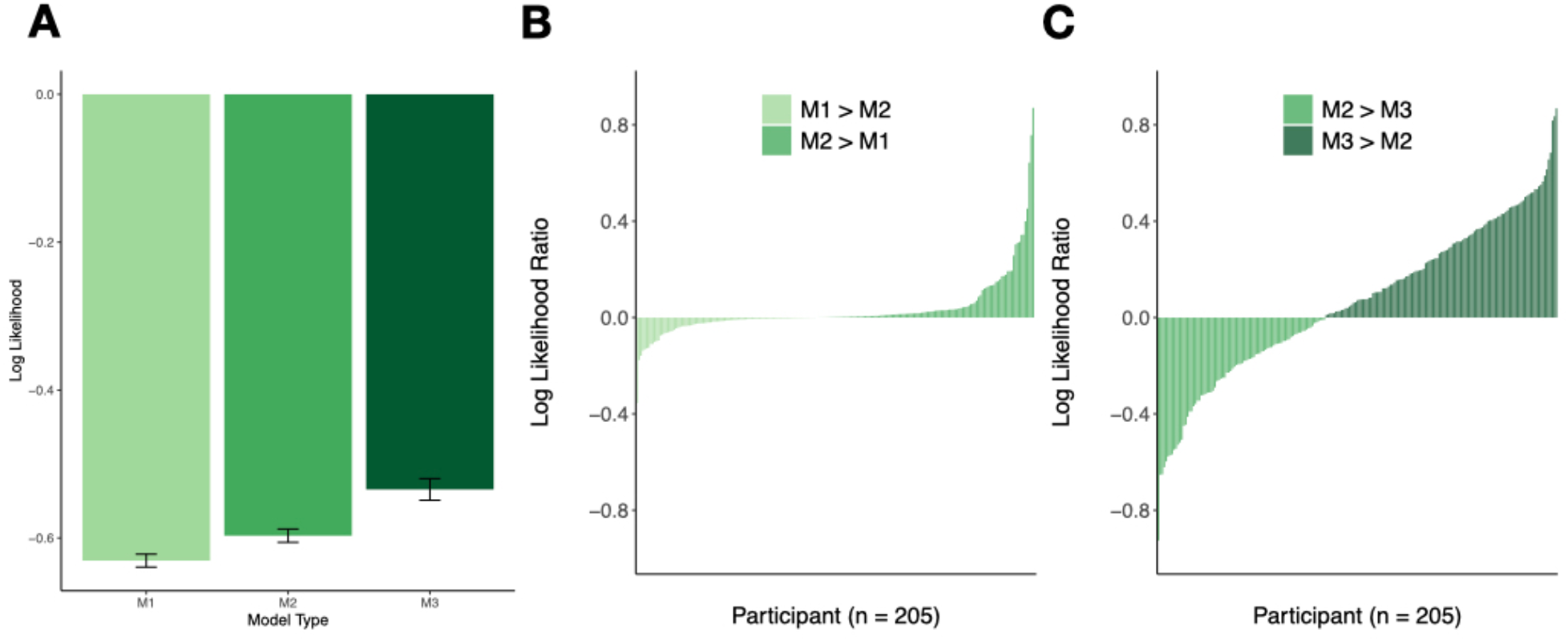
The reinforcement learning model with dual learning rates and stickiness is more predictive out of sample. **(A)** Mean log-likelihoods for held-out Block 4 data across all 205 participants for each of the three candidate models. Value closer to zero indicate higher predictive accuracy. Error bars reflect within-subject differences based on the method described in Cousineau (2015). **(B)** Plot of the difference in log likelihoods (model with one learning rate (M1) - model with two learning rates (M2)) averaged across trials and MCMC samples for each subject, on held out Block 4 data. Positive values indicate that M2 has greater predictive accuracy. A paired t-test shows that held-out log likelihoods are significantly higher on average for M2 (t(204) = 3.36; mean = 0.034 [0.014; 0.053]; p = 0.0009). **(C)** Plot of the difference in log likelihoods (two learning rate model (M2) - model with two learnings rates and stickiness (M3)) averaged across trials and MCMC samples, for each subject on held out Block 4 data. Positive values indicate that M3 has greater predictive accuracy. A paired t-test shows that held-out log likelihoods are significantly higher on average for M3 (t(204)= 2.76; mean = 0.063 [0.018; 0.11]; p = 0.0063).

Replicating the result reported in Gershman, 2016, we find that the more complicated of our three models better predicts out-of-sample data. Consequently, we will focus on this version of the model (M3) when comparing different modeling approaches. The marginal posterior distributions for the parameters of the group-level variables derived from M3 can be found in Supplemental Figure 2A. We have also plotted the empirical priors derived from the expectations of those group-level parameters in Supplemental Figure 2B.

## Evaluation of Model Fitting Methods

So far, we have used a hierarchical Bayesian approach to fit and compare reinforcement learning models. However, we have not yet shown that the hierarchical approach itself is any more useful than its various alternatives. To this end, we now turn our attention towards identifying the model-fitting technique that, when applied to M3, yields the most reliable results. We will do so by comparing the performance of 4 different model-fitting techniques that vary in whether and how they utilize group-level distributions to regularize individual estimates. The first two models represent opposite extremes: one avoids all group-level regularization to allow for infinite subject-specific variability (no pooling of information between subject-specific parameters), and the other allows for no subject-specific variability whatsoever (full pooling). The third and fourth models both constrain individual estimates based on empirical group-level priors, but while the former derives these priors from held-out data, the fully hierarchical model extracts and imposes group-level priors from a single data set. While we will focus on M3, the comparisons reported below are qualitatively similar when applied to all three models (see Supplement Figure 3 for comparison across model fitting techniques for M1 and M2).

### 1. Importance of empirical priors

Unless we can demonstrate that introducing group-level priors to constrain individual estimates improves predictive accuracy, the question of how to best derive them is moot.

To address this question, we first compared performance across two models – one that constrained individual estimates based on the group (described in Section 1: **Model**) and another in which these individual estimates were not pooled across subjects. We evaluated a Q-learning model with dual learning rates and stickiness (M3) on all 205 participants for both model-fitting techniques. For the no pooling model, this required adapting our hierarchical approach to eliminate group-level priors, and instead setting weakly informative hyperpriors on subject-specific parameters. Accordingly, the hyperpriors on individual learning rates (contained in the *α*_*n*_ vector), inverse temperatures, intercepts, and stickiness (contained in the *β*_*n*_ matrix) were specified as follows:

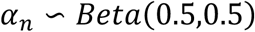

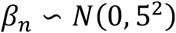

This approach essentially fits separate reinforcement learning models to each subject, and only constrains these estimates with weakly informative hyperprior distributions. It is perhaps the most common method for fitting reinforcement learning models in the literature (Davidow et al., 2016; Daw, 2011; Dezfouli & Balleine, 2013; Niv et al., 2015, 2012).

In order to evaluate the accuracy of the parameter estimates obtained from each of our two models of interest, we fit both of them on 3 out of 4 experimental blocks for each participant and computed the average log-likelihood of observations in the held-out block. The results are reported as the average log-likelihood for an observation for each participant, averaged across trials and posterior samples. As shown in **Figure 4A**, the full hierarchical model results in higher expected log likelihood for the held-out data (t(204) = 6.55; mean = 0.057 [0.040; 0.074]; p < 0.00005). This finding indicates that generalization is improved when group-level estimates are included as constraints on the extent of individual variability.

**Figure 4:**
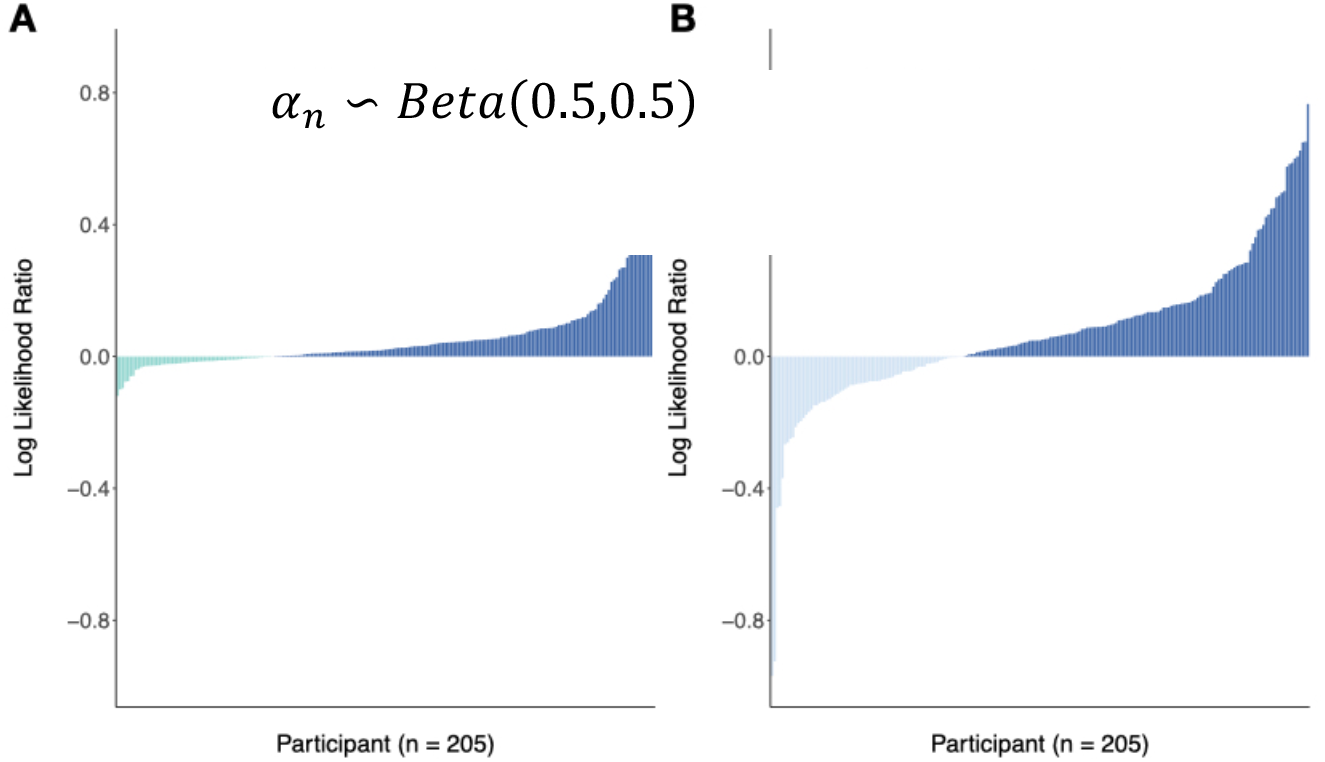
Hierarchical Bayesian models outperform two common alternatives. **(A)** Plot of the difference in log likelihoods (hierarchical model - no pooling model) averaged across trials and MCMC samples for each subject, on held out Block 4 data. Positive values indicate that the hierarchical model has greater predictive accuracy. A paired t-test indicates that held-out log likelihoods are significantly higher on average for the fully hierarchical model, meaning that group-level priors lead to greater predictive accuracy on held-out data (t(204) = 6.55; mean = 0.057 [0.040; 0.074]; p < 0.00005).). **(B)** Plot of the difference in log likelihoods (hierarchical model - full pooling model) averaged across trials and MCMC samples, for each subject on held out Block 4 data. Positive values indicate that the hierarchical model has greater predictive accuracy. A paired t-test shows that held-out log likelihoods are significantly higher on average for the fully hierarchical model (t(204) = 4.47, mean = 0.070 [0.040, 0.10], p < 0.00005).

Crucially, this leaves open the question of whether estimates should be allowed to vary across individuals at all. To answer this, we repeated the same model comparison procedure replacing the no pooling model with one that pools individuals completely, yielding only group-level estimates, which are used for each subject. We found that once again, the full hierarchical model results in higher expected log likelihood for held-out data (t(204) = 4.47, mean = 0.070 [0.040, 0.10], p < 0.00005, **Figure 4B**). This further supports the notion that a model’s predictive accuracy is improved by accounting for individual variability while constraining it, which is the premise underlying most multi-level modeling (Gelman & Hill, 2007). Empirical priors seem to provide the right balance between too much and too little group-level influence for reinforcement learning models. A crucial question, then, lies in which is the best strategy for deriving them.

### 2. Derivation of empirical priors

While we have shown that constraining individual estimates based on data-derived priors leads to improved predictive accuracy, the question still remains of how these empirical priors should be obtained. In Gershman (2016), the data from 165 out of 205 participants are set aside for the purpose of generating reliable group-level priors. This involves fitting a reinforcement learning model with weakly informative priors on the 165-person subset and once subject-specific parameter estimates are obtained, estimating group-level priors by moment matching. The resulting empirical priors are then applied to the remaining 40 participants, which leads to improved model fit and generalizability for this subset of participants.

Using the same dataset, we replicated the approach by fitting the version of our model that includes only weakly informative group-level priors on 165 of the participants. Following the method described above, we then used a moment-matching function to approximate the parameter estimates that best describe the distributions of subject-level parameters derived from the models fit to each subject. More specifically, this moment-matching function uses gradient descent on the distribution of individual MAP estimates to derive the best parameters for a group-level probability distribution. The outputs of the function (**Table 1**) describe the distributions of the group-level priors that we then used on a separate dataset, consisting of the 40 remaining participants.

**Table 1:** Empirical Prior Distributions.

| Learning Rate | Intercept | Inverse Temperature | Stickiness |
| --- | --- | --- | --- |
| $\alpha_1 \sim \text{Beta}(1.15, 1.84)$<br>$\alpha_2 \sim \text{Beta}(1.01, 1.20)$ | $\beta_0 \sim \text{Normal}(0.050, 0.63^2)$ | $\beta_1 \sim \text{Normal}(2.55, 3.18^2)$ | $\beta_2 \sim \text{Normal}(-0.13, 2.8^2)$ |

We first replicate the results from Gershman, 2016: including out-of-sample priors improves predictive accuracy when compared to a model run on the same 40 participants without any group-level constraints (the same “no pooling” model used for comparison in the previous section) (t(39) = 3.18; mean = 0.039 [0.014; 0.065]; p = 0.0029, **Figure 5 A&B**). The inclusion of out-of-sample priors also yields a higher average predictive accuracy on held-out data than the full pooling model that ignores individual variability entirely (t(39) = 2.24; mean = 0.077 [0.0074; 0.15]; p = 0.031, **Figure 5A**).

**Figure 5:**
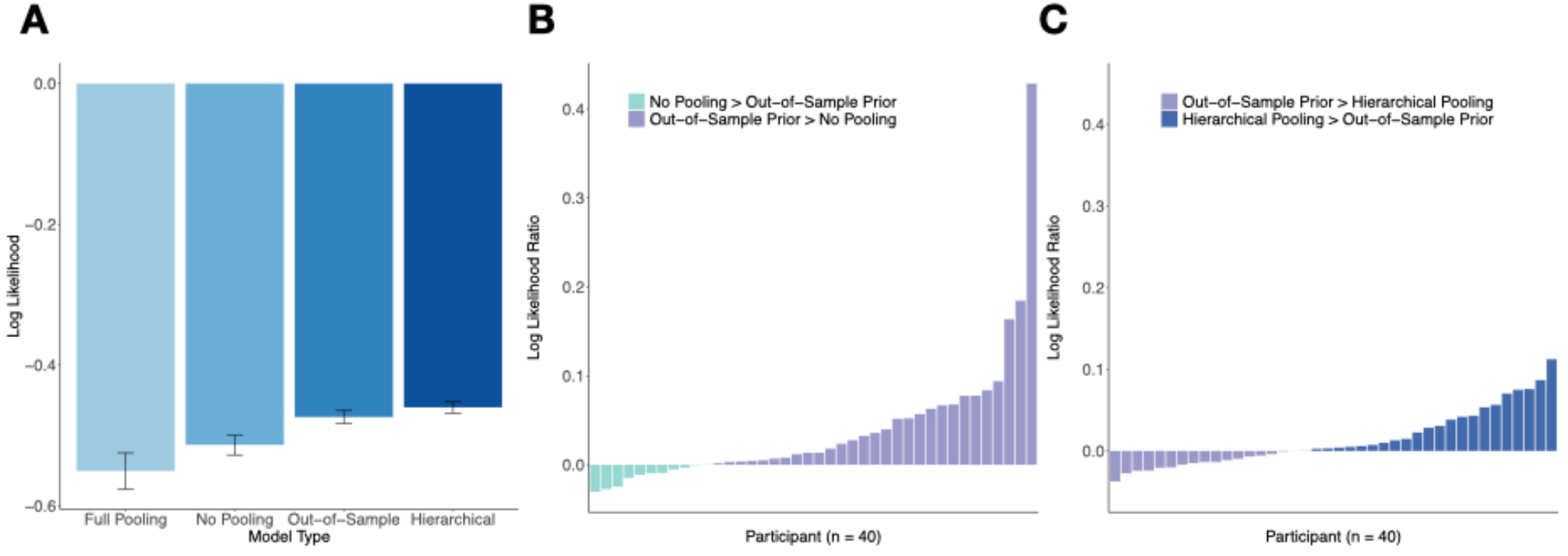
Hierarchical Bayesian modeling has higher predictive accuracy than alternative approaches. **(A)** Mean of the log-likelihoods for held-out Block 4 data across 40 participants for each of the four candidate models. Value closer to zero indicate higher predictive accuracy. Error bars reflect within-subject differences based on the method described in Cousineau (2015). **(B)** Plot of the difference in log likelihoods (model with out-of-sample priors - no pooling model) averaged across trials and MCMC samples for each subject, on held out Block 4 data. Positive values indicate that the model that uses out-of-sample empirical priors has greater predictive accuracy. A paired t-test shows that held-out log likelihoods are significantly higher on average for the model with out-of-sample priors (t(39) = 3.18; mean = 0.039 [0.014; 0.065); p = 0.0029). **(C)** Plot of the difference in log likelihoods (hierarchical model - model with out-of-sample priors) averaged across trials and MCMC samples, for each subject on held out Block 4 data. Positive values indicate that the full hierarchical model has greater predictive accuracy. A paired t-test shows that held-out log likelihoods are significantly higher on average for the hierarchical model, meaning that hierarchically enforcing group-level priors leads to greater predictive accuracy than extracting the priors from held-out data (t(39)= 2.48; mean = 0.014 [0.0025; 0.025); p = 0.018).

The question remains, however, of whether it is necessary to derive these priors from a separate, held-out group of participants. One of the benefits of the hierarchical Bayesian approach is that it generates group-level priors and individual estimates using the same data. This method circumvents the need to discard data for the purpose of generating reliable empirical priors because it accomplishes both tasks at the same time. Furthermore, because of the two-way nature of hierarchical modeling – group estimates affect individual estimates and vice versa – the parameter values for the participants that would normally be discarded will also be more accurate. For these held-out participants, this is the equivalent of comparing the no pooling approach to the empirical pooling approach, which we have already shown increases predictive accuracy (**Figure 5, A&B)**.

To assess the computational benefits of extracting empirical priors from a held-out group of participants, we can compare the log-likelihood estimates from this model to those of the fully hierarchical approach. **Figure 5C** shows that log-likelihood estimates for Block 4 data are in fact higher when the group-level priors are derived and fit simultaneously using a hierarchical Bayesian approach on just 40 participants, compared to when the priors are derived from a separate group of 165 people (t(39)= 2.48; mean = 0.014 [0.0025; 0.025]; p = 0.018). Thus, the hierarchical model allows for greater predictive accuracy, while also cutting down the number of participants needed to fit the model by over 80% (40 participants instead of 205).

### 3. Individual differences

Given that hierarchical pooling constrains individual estimates based on group-level priors, one possible concern is that the approach might reduce our ability to detect real individual differences. To assess this concern, we can compare how well individual differences in task performance track with related parameter estimates across model-fitting techniques. Specifically, we can measure the correlation between each subject’s performance on the task – operationalized as mean-squared prediction error – and inverse temperature. We would expect that these two values should be correlated, with higher sensitivity to reward leading to lower error (Gershman, 2016). Because the full pooling model does not allow for individualized parameter estimates, we have excluded this approach from the comparison. Across the three other model-fitting techniques, we see a negative correlation between mean-squared error and inverse temperature, as expected (Figure 6). Importantly, the magnitude of the correlation is comparable for both hierarchical pooling and out-of-sample priors. Thus, hierarchical priors do not in principle limit our ability to detect individual differences.

**Figure 6:**
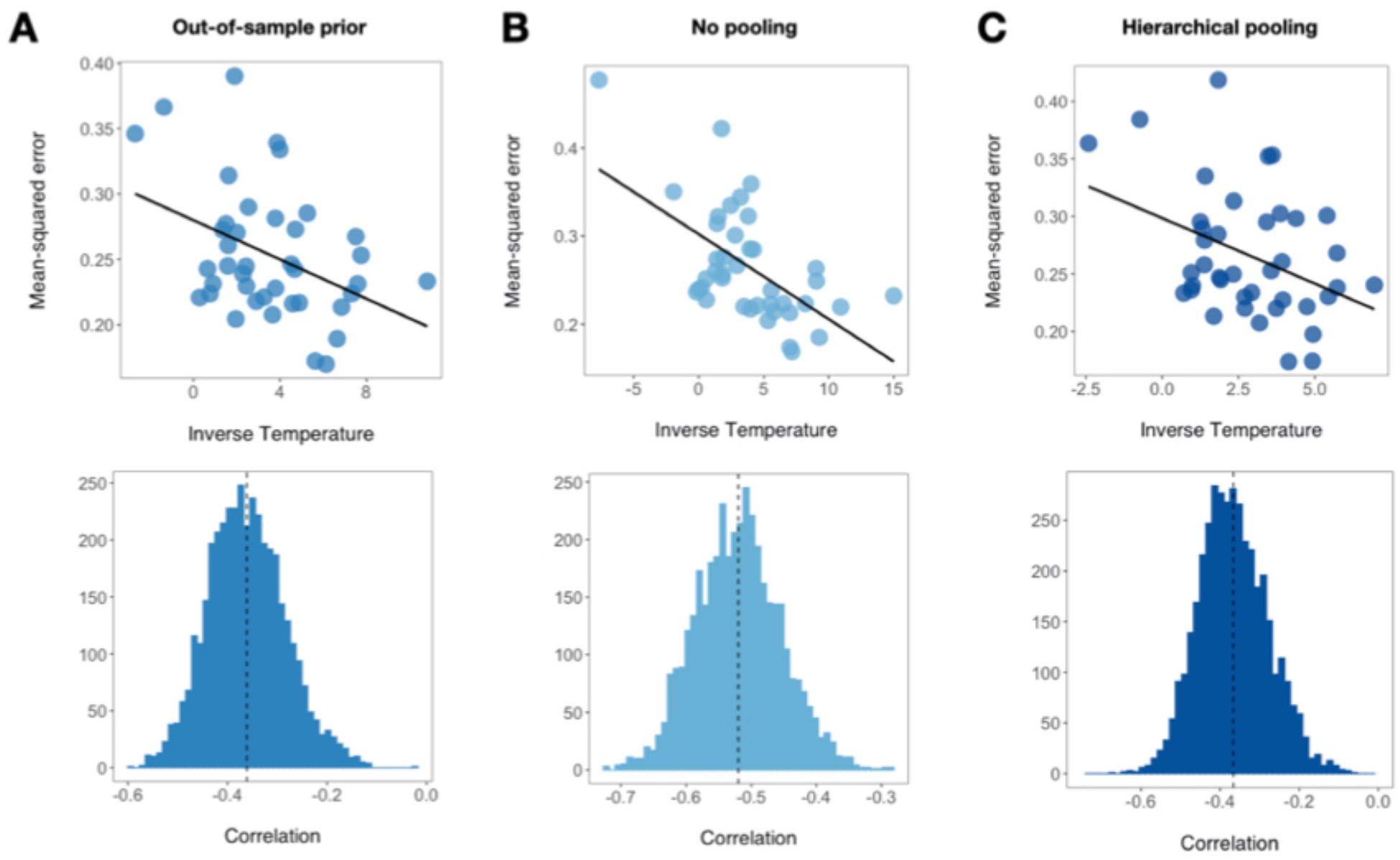
The use of hierarchical priors preserves sensitivity to individual differences. **Top Row:** Correlation between reward prediction accuracy (operationalized as mean-squared model-derived PE) and inverse temperature in 40 participants fit using **(A)** out-of-sample priors, **(B)** with no pooling, and **(C)** using hierarchical priors. Each point on the scatter plot corresponds to the expected value of a subject’s posterior distribution, and black lines are fits the regression of inverse temperate on mean-squared prediction error. **Bottom Row:** Posterior histograms of the correlation for models fit with **(A)** out-of-sample priors, **(B)** no pooling, and **(C)** hierarchical priors. For each sample from the posterior, we computed the correlation between subject-level inverse temperature and mean-squared prediction error. The process was repeated for every sample sample to generate a posterior distributions over correlations.

## Discussion

In this paper, we have provided an overview of the implementation and benefits of hierarchical Bayesian models of reinforcement learning. We began with a tutorial on how to fit such models, ranging from potential parameters to include to how and when to incorporate priors. We found that fitting all three versions of the reinforcement learning model hierarchically, predictive accuracy was highest for the model that included dual learning rates as well as a stickiness parameter (M3). In the subsequent section, we focused on M3 specifically and compared its performance across four different model-fitting techniques.

In line with previous work (Daw, 2011; Efron, 1996; Gershman, 2016), we have argued that data-driven group-level priors improve reinforcement learning models in several ways. Their inclusion constitutes a reliable middle-ground in the variance-bias tradeoff, as it allows for subject-specific estimates and individual differences, while also constraining these estimates based on the group. This approach improves predictive accuracy, as it pushes very uncertain parameters towards a group average. The improvements group-level priors provide become evident when we compare the predictive accuracy of reinforcement learning models with no group-level constraints to those with empirical priors.

Furthermore, we have compared the hierarchical model’s performance to that of a model that also leverages empirical priors, but derives them from a separate dataset. Doing so reveals that the benefits of the hierarchical model are two-fold: not only does it predict held-out data with greater accuracy, it does so with significantly fewer data points and, consequently, is more efficient when considered in an experimental context. As data collection takes time and slows scientific progress, we see this as an important virtue of the hierarchical approach.

Although it is not the main focus of the paper, we believe that Bayesian models of reinforcement learning also provide a more transparent handling of uncertainty than methods that rely on approximating point estimates. They do so by yielding a full distribution of possible values for all parameters, which avoids the potential issues caused by selecting just one value through maximum likelihood estimation. The fully Bayesian hierarchical model we have described maintains a measure of epistemic uncertainty (the uncertainty derived from trying to map observations onto parameters) throughout, acknowledging the ambiguity inherent in computational modeling at each level of analysis.

There are several potential limitations to our methodology. To begin with, we have limited our model comparisons to a small number of relatively straightforward reinforcement learning models in order to focus on a detailed illustration of the hierarchical approach. Insofar as the risks of overfitting increase with the effective number of parameters, however, it is possible that more complex RL models that include more than just our maximum of five parameters (M3) would yield different results. Nonetheless, to our knowledge, the possibility that out-of-sample priors might perform better than hierarchical pooling for more complex models has not been demonstrated.

Secondly, model performance is inherently tied to the metric used to assess it. Throughout this paper, we have computed or estimated log-likelihoods on held-out data for each subject in order to compare one model fitting technique to the next. While this method is valid and echoes more standard information criteria, it does not predict observations from new subjects. Relatedly, one comparative advantage of deriving empirical priors from held-out data is that the estimated group distribution is derived from a group of participants that is kept separate. Thus, it is possible that this distribution would generalize better to new people, as it is more removed from the in-sample group. Future work should explore the effect of hierarchical modeling decisions on prediction to new subjects and experimental contexts.

One final caveat to emphasize is that hierarchical Bayesian models do not constitute a one-size-fits-all solution for any theoretical framework. When deciding how to fit a statistical model on a given dataset, theory should take precedence over statistical concerns, and if the model is inadequate, any specific inference technique will be of little use. Our goal here is only to show that for a very restricted but widely applied set of reinforcement learning models (Q-learning models), the hierarchical Bayesian approach can be beneficial.

Individual difference measures play a crucial role in computational modeling in psychology and neuroscience. Latent parameters from reinforcement learning models, for example, are often correlated with other behaviors and demographic variables, or even with clinical traits in psychiatric populations (Huys et al., 2016; Maia & Frank, 2011; Radulescu et al., 2016; Rouhani & Niv, 2019). In the field of cognitive neuroscience, parameter estimates are also often used to track correlates of brain activity, such as BOLD activation (Behrens et al., 2007; Cohen et al., 2017; McClure et al., 2003; Niv, 2009; O’Doherty et al., 2007; O’Reilly, 2013). Given the prevalence of these methods, the extraction of reliable parameter estimates is crucial: the strength of a study’s conclusions is contingent on the stability of the estimated parameters (but see R. C. Wilson & Niv, 2015 for an interesting counterpoint). In recognition of this delicate situation, hierarchical models, especially the Bayesian models we describe here, provide an effective method for generating reliable estimates with appropriate levels of uncertainty.

## Acknowledgments

The authors thank Matti Vuorre, Daphna Shohamy, Samuel Lippl, and Jonathan Nicholas for useful discussion and helpful comments on a previous draft of the paper. This research did not receive any specific grant from funding agencies in the public, commercial, or not-for-profit sectors.

