## Supplemental Figures for "Hierarchical Bayesian Models of Reinforcement Learning: Introduction and comparison to alternative methods"

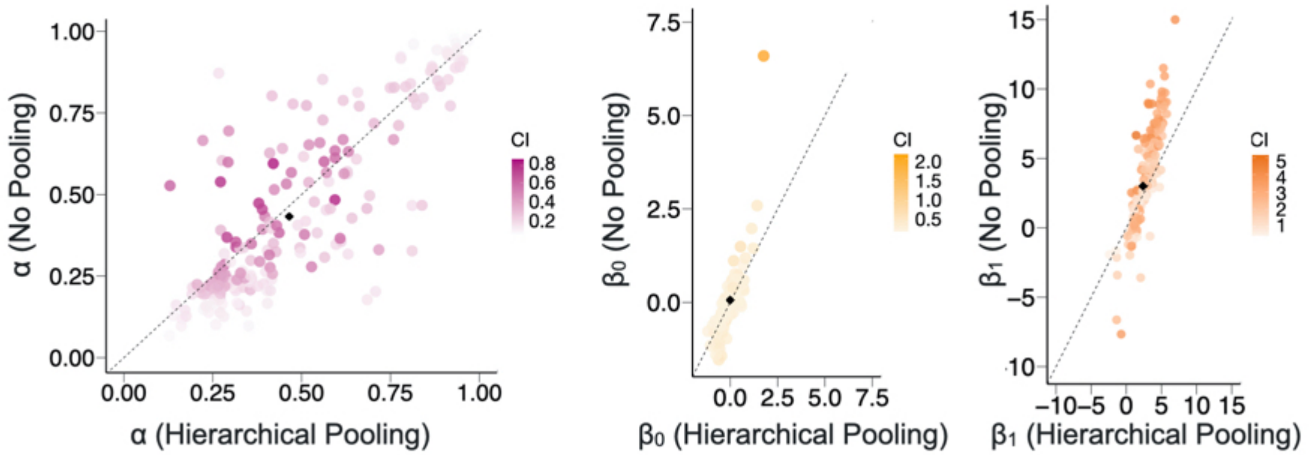


**Figure S1: The use of hierarchical priors helps regularize parameter estimates of initially uncertain or extreme parameters.** Plots of subject-level parameter estimates calculated using the expected value of each subject's posterior distribution from the no pooling vs. hierarchical pooling models. For unconstrained parameters $\beta_0$ and $\beta_1$, hierarchical priors shrink extreme parameter values towards the group mean. The effect is less clear for α. Shading corresponds to the width of the confidence interval around each subject's parameter estimate in the no pooling model, taken as an index of uncertainty.

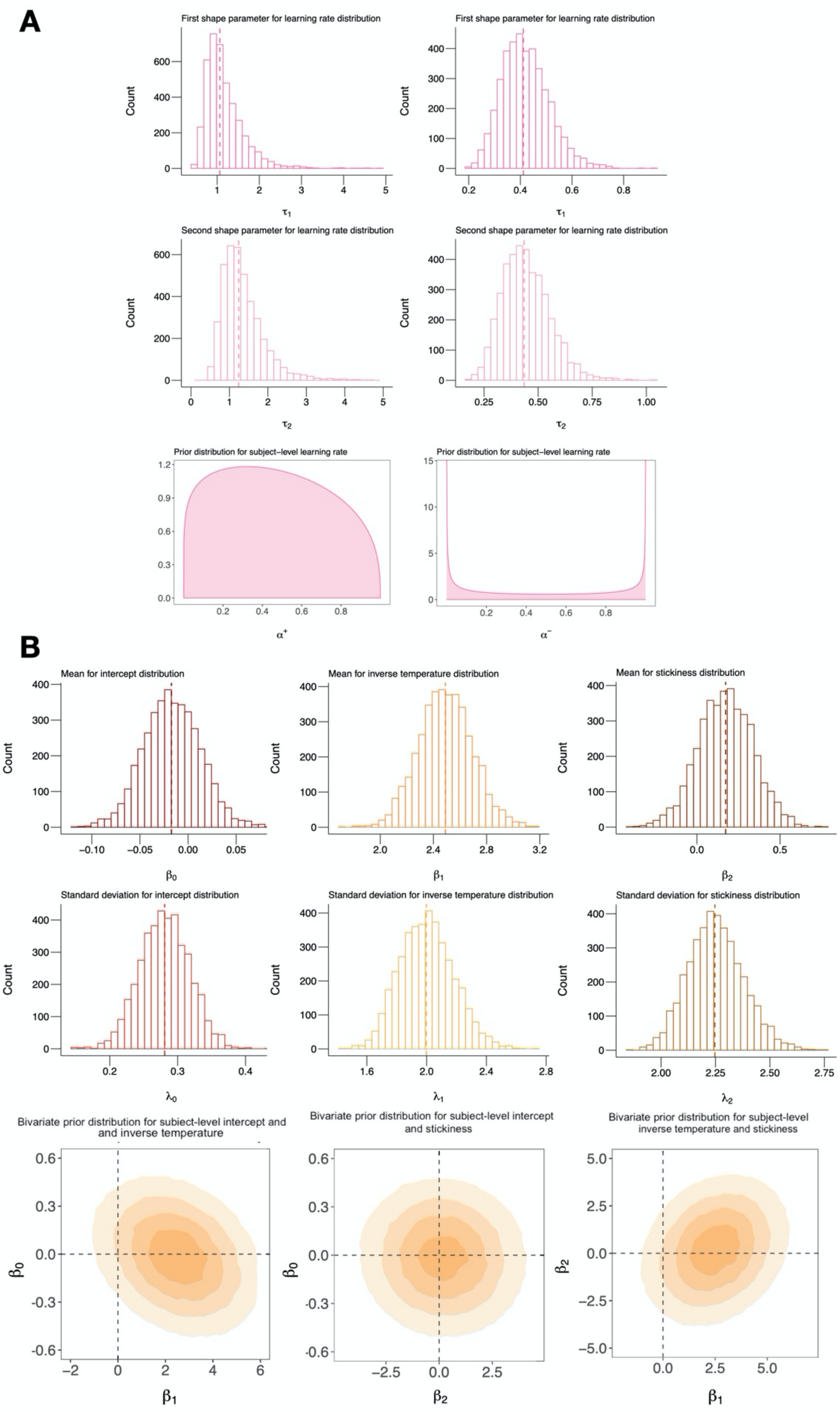


**Figure S2: Posterior histograms for group-level parameters and expected empirical prior distributions in model M3. (A) Top:** Posterior histograms for group-level parameters on learning rate estimates, fit hierarchically on model M3. Dashed line corresponds to the median of the each distribution. **Bottom:** Expected empirical prior distribution for learning rates in model M3. While the prior distribution for $\alpha^+$ provides support for incremental learning — the most likely values of $\alpha^+$ are between 0.2 and 0.6 — the prior distribution for $\alpha^-$ highlights that participants may respond to negative outcomes either by ignoring them ($\alpha^- = 0$) or using a win-stay-lose-shift strategy ($\alpha^- = 1$). **(B) Top:** Posterior histograms for group-level parameters on intercept, inverse temperature, and stickiness estimates, fit hierarchically on model M3. **Bottom:** Prior distributions for intercept, inverse temperature, and stickiness, parameterized using the mean of the posteriors. Each plot is multivariate Normal, as it includes model-derived covariance estimates for each combination of parameters ($\beta_0$ and $\beta_1$; $\beta_0$ and $\beta_2$ ; $\beta_1$ and$\beta_2$).

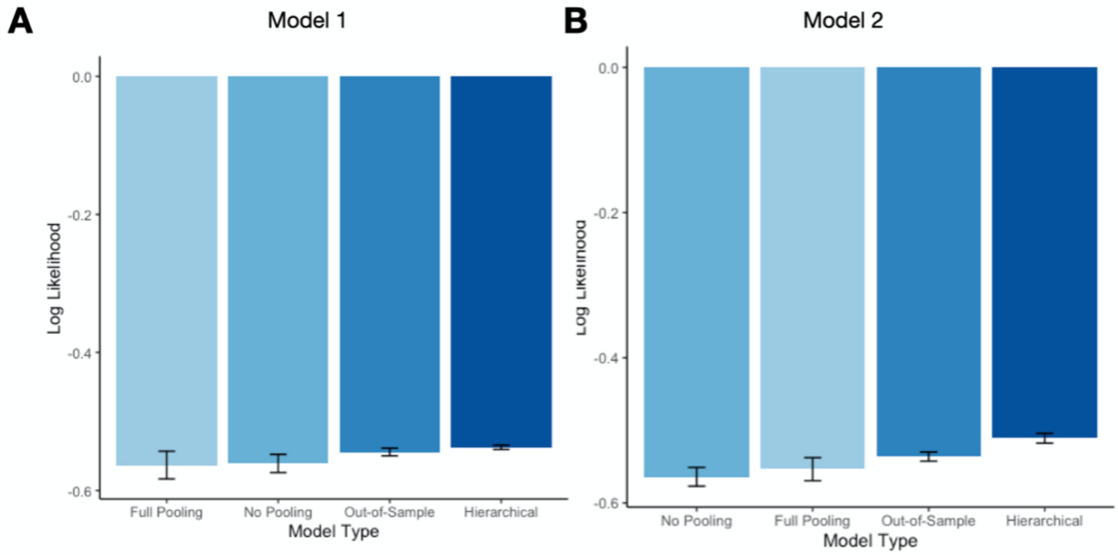


**Figure S3: Hierarchical pooling improves predictive accuracy for M1 and M2.** Mean of the log-likelihoods for held-out Block 4 data across 40 participants for each of the four candidate model-fitting techniques. Value closer to zero indicate higher predictive accuracy. Error bars reflect within-subject differences based on the method described in Cousineau (2015).
